# Subregions of DLPFC display graded yet distinct structural and functional connectivity

**DOI:** 10.1101/2021.06.14.448404

**Authors:** JeYoung Jung, Matthew A. Lambon Ralph, Rebecca L. Jackson

## Abstract

The human dorsolateral prefrontal cortex (DLPFC, approximately corresponding to Brodmann areas 9 and 46) has demonstrable roles in diverse executive functions such as working memory, cognitive flexibility, planning, inhibition, and abstract reasoning. However, it remains unclear whether this is the result of one functionally homogeneous region or whether there are functional subdivisions within the DLPFC. Here, we divided the DLPFC into seven areas along with rostral-caudal and dorsal-ventral axes anatomically and explored their respective patterns of structural and functional connectivity. *In vivo* probabilistic tractography and resting-state functional magnetic resonance imaging were employed to map out the patterns of connectivity from each DLPFC subregions. Structural connectivity demonstrated graded intra-regional connectivity within the DLPFC. The patterns of structural connectivity between the DLPFC subregions and other cortical areas revealed that he dorsal-rostral subregions was restricted to connect to other frontal and limbic areas, whereas the ventral-caudal region was widely connected to frontal, temporal, parietal, and limbic cortex. Functional connectivity analysis demonstrated that subregions of DLPFC were strongly interconnected to each other. The dorsal subregions were associated with the default mode network (DMN), while middle dorsal-rostral subregions were linked with the multiple demand network (MDN), respectively. Similar to the results of structural connectivity, the ventral-caudal subregion showed increased functional coupling with both DMN and MDN. Our results suggest that DLPFC may be subdivided by the diagonal axis of the dorsal-ventral axis and rostral-caudal axis, which support the patterns of connectivity the parts of the DLPFC reflects its integrative executive function.

## Introduction

The dorsolateral prefrontal cortex (DLPFC) approximately corresponds to Brodmann areas (BA) 9 and 46 and consists of the lateral part of superior frontal gyrus (SFG) and middle frontal gyrus (MFG) (Brodmann, 1908; Walker, 1940; Petrides and Pandya, 1999). More recently, Petrides (2005b) designated DLPFC as BA 9, BA 46, and BA 9/46. Previous anatomical and functional studies have demonstrated differences in the subparts of lateral prefrontal cortex connectivity including the DLPFC (For the reivew, see Petrides, 2005b; Thiebaut de Schotten et al., 2012; Cieslik et al., 2013). Although the DLPFC can be divided into two or three subregions cytoarchitectonically, the functional role of each subregion is not clear.

DLPFC plays an important role in executive functions, such as working memory, cognitive flexibility, planning, inhibition, and abstract reasoning (Miller and Cummings, 2007) and is connected to a variety of brain regions including the thalamus, basal ganglia, hippocampus, and associative cortex such as posterior temporal, and parietal areas (Petrides and Pandya, 1984; Morris et al., 1999; Petrides, 2005b; Yeterian et al., 2012). Anatomical studies have demonstrated that different areas within the DLPFC receive their input from distinct subparts of the parietal cortex (Petrides and Pandya, 1984; Thiebaut de Schotten et al., 2012) and that the functional role of the DLPFC may be partly determined by its anatomical connections to other brain regions (Morecraft et al., 2004; Hoshi, 2006). Strong evidence from studies with both human and primates suggests an anterior-posterior axis of functional organization of the lateral prefrontal cortex (Koechlin et al., 2003; Petrides, 2005b, a; Thiebaut de Schotten et al., 2012). Lesions in the caudal DLPFC (BA 46 and 9/46) are associated with a deficit on monitoring of information in working memory whereas the further caudal lesions in BA6 and 8 impair tasks requiring selection between alternative choices (Petrides, 1985, 2005a). The rostral mid-lateral prefrontal regions (BA 10 and 46) play a more abstract role in cognitive control (Petrides, 2005b; Moayedi et al., 2015). In addition to the caudal-rostral axis, there is a dorsal-ventral axis of organization of the mid-lateral prefrontal cortex (DLPFC: BA 46 and 9/46) (Petrides and Pandya, 1999; Petrides, 2005b). In monkey, the dorsal DLPFC (BA9/46d) plays a role in motor planning, multi-tasking, and maintaining goals whereas the ventral DLPFC (BA9/46v) is preferentially involved in the visuospatial information of attended signals and cues. These findings suggest a possibility that some functionally distinct subparts may exist within the DLPFC.

Despite the well-documented role of the DLPFC in various executive functions, it remains unclear whether functional subdivisions are present within the DLPFC. Although numerous fMRI studies report DLPFC activation, the exact location and extent of activation sites vary according to tasks used in those studies (Nee et al., 2007; Zheng et al., 2008; Rottschy et al., 2012; Bahlmann et al., 2014; Kohn et al., 2014). One study parcellated the DLPFC using a meta-analysis of task-dependent and task-independent connectivity (Cieslik et al., 2013). They delineated the DLPFC into two regions with different connections: the anterior subregion co-activated with the ACC and the posterior subregion co-activated with the parietal cortex. These findings support functional variation along a rostral-caudal axis in the DLPFC (Petrides, 2005a). Together, the functional heterogeneity and diversity of anatomical connections in the human DLPFC suggests it may consist of functionally distinct subregions.

Here, we test whether there are connectivity differences across the rostral-caudal and dorsal-ventral axes in the DLPFC that would lead to different functional subregions. We used diffusion weighted imaging (DWI) and resting state fMRI to explore the structural and functional connectivity across the DLPFC in human participants. To improve signal in ventromedial frontal and anterior temporal regions, while maintaining signal across the whole brain, we employed probabilistic tractography of distortion-corrected DWI (Embleton et al., 2010) and seed-based analysis of dual-echo resting state fMRI (Halai et al., 2014). The DLPFC was separated into seven different areas based on the rostral-caudal and dorsal-ventral divisions of the Brodmann regions (BA9, 46, and 9/46) (Petrides, 2005b). For each seed region, we investigated and compared the structural connectivity to sixty-three target regions including left frontal, temporal, parietal, and limbic cortex and the whole brain functional connectivity. We hypothesized that there would be functional subregions of the DLPFC determined by their structural or functional connectivity and that there would be differential patterns of structural and functional connectivity within the DLPFC along the rostral-caudal or dorsal-ventral axis.

## Materials and Methods

### Subjects and data acquisition

Two different datasets were employed in this study. Dataset 1 included 24 healthy, right-handed subjects (11 females; mean age = 25.9, range 19-47), whereas dataset 2 included 78 healthy right-handed subjects (57 females; means age = 25.2, range 20-44). Each dataset has been utilized for various investigations: Dataset 1 (Cloutman et al., 2012; Bajada et al., 2016; Bajada et al., 2017; Jung et al., 2017; Jackson et al., 2020) Dataset 2 (Jackson et al., 2016; Jackson et al., 2018; Jung et al., 2018; Jackson et al., 2020). Dataset 1 consisted of distortion-corrected DWI and structural MR imaging. DWI was performed using a pulsed gradient spin echo-planar sequence, with TE = 59ms, TR ≈ 11884ms, G = 62 mTm^−1^, half scan factor = 0.679, 112 × 112 image matrix reconstructed to 128 × 128 using zero padding, reconstructed resolution 1.875 × 1.875 mm, slice thickness 2.1 mm, 60 contiguous slices, 61 non-collinear diffusion sensitization directions at b = 1200 smm^−2^ (Δ = 29.8 ms, δ = 13.1 ms), 1 at b = 0, SENSE acceleration factor = 2.5. Acquisitions were cardiac gated using a peripheral pulse unit positioned over the participants’ index finger or an electrocardiograph. For each gradient direction, two separate volumes were obtained with opposite polarity *k*-space traversal with phase encoding in the left-right/right-left direction to be used in the signal distortion correction procedure (Embleton et al., 2010). A co-localized T2 weighted turbo spin echo scan, with in-plane resolution of 0.94 × 0.94 mm and slice thickness 2.1 mm, was obtained as a structural reference scan to provide a qualitative indication of distortion correction accuracy. A high resolution T1-weighted 3D turbo field echo inversion recovery image (TR ≈ 2000 ms, TE = 3.9 ms, TI = 1150 ms, flip angle 8º, 256 × 205 matrix reconstructed to 256 × 256, reconstructed resolution 0.938 × 0.938 mm, slice thickness 0.9 mm, 160 slices, SENSE factor = 2.5) was used.

Dataset 2 included resting-state fMRI and structural MR imaging. To cover the whole brain without signal dropout around the rostral temporal and inferior frontal cortices, a dual-echo fMRI protocol was performed (Poser et al., 2006; Halai et al., 2014). This involves parallel acquisition at a short echo (12ms) leading to less signal loss in areas of high magnetic susceptibility and a standard long echo (35ms) to maintain high contrast sensitivity throughout the brain. The results from the 2 echoes were combined using linear summation, previously shown to be optimal. The fMRI parameters included 42 slices, 80 × 80 matrix, 240× 240 × 126mm FOV, in-plane resolution 3 × 3, slice thickness 4mm. 130 volumes were collected over 6.25 minutes. T1-weighted structural images were acquired using a 3D MPRAGE pulse sequence with 200 slices, in-planed resolution 0.94 × 0.94m slice thickness 1.2mm, TR = 8.4ms, TE =3.9ms. During resting-state fMRI, all subjects were instructed to keep their eyes open and look at the fixation cross. Imaging data were acquired on a 3T Philips Achieva scanner (Philips Medical System, Best Netherlands). The study was approved by the local ethics committee and all participants provided written informed consent forms.

### Definition of seed regions and target masks

In order to divide the DLPFC (BA 9 and BA46) on the rostral-caudal dimension and to explore differences in DLPFC connectivity, 7 anatomically defined regions of interest (ROIs) were defined in the left hemisphere (Fig.1A): two located on BA 9 (anterior: 9a, posterior: 9p), two placed in dorsal-middle frontal gyrus (anterior: 9/46da, posterior: 9/46dp), one was on BA 46 (46), and two placed in ventral-middle frontal gyrus (anterior: 9/46va, posterior: 9/46vp). The seeds regions were identified as a sphere with 6 mm radius in the MNI template brain based on topographic description and defined carefully without overlapping each other. 63 target regions covering frontal, temporal, parietal, and limbic cortex were defined using WFU Pick Atlas (Maldjian et al., 2003) and SPM Anatomy toolbox (Eickhoff et al., 2005). It should be noted that the occipital lobe was not included in this study because there was no direct white matter pathways connecting the DLPFC and occipital lobe (Petrides and Pandya, 1984; Morris et al., 1999; Petrides, 2005b; Yeterian et al., 2012). The frontal lobe regions included BA 10 (frontal pole: FP), BA 44 (pars opercularis), BA 45 (pars triangularis), BA47 (pars orbitalis), medial orbitofrontal cortex (medOFC), lateral OFC (latOFC), supplementary motor area (SMA), and primary motor cortex (M1). The parietal lobe had 7 regions in superior parietal cortex (SPC: 5L, 5M, 5Ci, 7A, 7PC, 7M, 7P), 3 regions in intraparietal sulcus (IPS: IPS1, IPS2, IPS3), and 7 regions in inferior parietal cortex (IPC: PFop, PFt, PF, PFm, PFcm, PGa, PGp). The temporal lobe had 20 ROIs covering superior temporal gyrus (STG), middle temporal gyrus (MTG), inferior temporal gyrus (ITG), fusiform gyrus (FG), parahippocampal gyrus (PhG) (please, see the detailed location of the temporal ROIs for Jung et al., 2017; Jung et al., 2018). The limbic lobe included insular, amygdala, hippocampus, caudate, putamen, pallidum, thalamus, and 3 regions of cingulate cortex (anterior cingulate cortex; ACC, middle CC; MCC, posterior CC; PCC). The DARTEL (diffeomorphic anatomical registration through an exponentiated lie algebra) toolbox (Ashburner, 2007) was used to transform the seeds and masks from the MNI space into each individual’s native diffusion space. The transform was estimated using each subject’s T1-weighted image coregistered to their diffusion weighted images. The accuracy of the transformation of ROIs into native space was inspected using the anatomical images. For resting-state functional connectivity analysis, ROIs without DARTEL transformations were used as analysis was performed in MNI space.

### Probabilistic tractography

Unconstrained probabilistic tractography was performed using the PICo software package (Parker and Alexander, 2005), sampling the orientation of probability density functions (PDFs) which was generated using constrained spherical deconvolution (Tournier et al., 2008) and model-based residual bootstrapping (Haroon et al., 2009; Jeurissen et al., 2011). 20,000 Monte Carlo streamlines were initiated from each voxel in the DLPFC seed regions. Step size was set to 0.5 mm. Stopping criteria for the streamlines included terminating if the pathway curvature over a voxel was greater than 180º, or the streamline reached a physical path limit of 500 mm. A single whole-brain probabilistic map was generated for each of the 7 seed ROIs for each participant. Probability maps were masked with each ROI and the maximum connectivity value (ranging from 0 to 20,000) between the seeds and each mask was extracted. The resultant connectivity matrices were subjected to a double threshold to ensure that only connections with high probability in the majority of participants were considered. For the first-level individual threshold, following the approach described by Cloutman et al. (2012), the λ-value of the Poisson distribution identified was used to determine a threshold value at p = 0.025. For the second-level group threshold, we used two criteria for consistency (over 75% of participants, i.e., at least 18/24 participants and over 50% of participants, i.e., at least 12/24 participants).

### Resting-state fMRI data analysis

Pre-processing was performed using SPM 8 and the data processing assistant for Resting State fMRI (DPARSF Advanced Edition, V2.3) toolbox. The first two volumes were discarded to allow for magnetic saturation effects. The images were slice-time corrected, realigned, and coregistered to the individual’s structural image using SPM 8. Censoring was applied using a threshold of greater than 3mm of translation or 1 degree of rotation, which resulted in the exclusion of 6 participants from further analysis. Within DPARSF nuisance covariates were regressed out and the images were normalised using DARTEL, smoothed with an 8mm full-width half maximum (FWHM) Gaussian kernel. The results were filtered at .01 - .08 Hz.

Nuisance covariates were regressed out including 24 motion parameters calculated from the 6 original motion parameters using Volterra expansion (Friston et al., 1996), which was shown to improve motion correction compared to the 6 parameters alone (Yan et al., 2013; Power et al., 2014). Additional covariates were included for outlier time points with a with a z-score greater than 2.5 from the mean global power or more than 1mm translation as identified using the ARtifact detection Tools software package (ART; www.nitrc.org/projects/artifact_detect). These were entered as covariates with white matter, CSF and global tissue signal. Then, linear detrending was performed. Seed-based functional connectivity analyses were performed using DPARSF. Functional connectivity maps from the seeds were z-score normalised. One sample t-tests were used to detect areas with significant connectivity to the seed regions. The resultant images were thresholded at p < 0.001, FWE-corrected at the cluster level. Comparisons between the functional connectivity maps of different seed regions were conducted using paired t-tests.

## Results

### Structural connectivity patterns across the DLPFC

Using probabilistic tractography, the structural connectivity for each DLPFC seed was identified, see Table 1 and Fig. 1. The full pattern of connectivity across the brain may be seen in Fig. 2. There was strong intra-DLPFC connectivity on the dorsal-ventral axis (along the gyri). Additionally, the mid-DLPFC regions (9/46da and 9/46dp) showed the strongest intra-DLPFC connectivity, with connections to more dorsal and ventral areas (Fig. 1B). Across the DLPFC, there was a high level of connectivity with limbic regions, especially the insular and basal ganglia (caudate, putamen, and pallidum) (Fig. 1C). The ventral-caudal seeds (9/46dp, 9/46va, and 9/46vp) showed structural connections with the thalamus. Only 9/46vp had a connection to hippocampus. However, no direct connection was identified between any seed regions and the amygdala. Fig. 1D shows the pattern of connectivity between the DLPFC seed regions and other lateral associative cortices. All DLPFC seed regions showed strong connectivity with the frontal pole (FP) and inferior frontal gyrus (IFG: BA44 and BA45) but not the most ventral aspects of the prefrontal cortex, including pars orbitalis (BA47) and the OFC. Only the ventral-caudal seeds (9/46va and 9/46vp) had strong evidence of connections to primary and supplementary motor regions. Additionally, only the 9/46vp seed connects to somatosensory and dorsal parietal regions (7PC and IPS). It should be noted that the DLPFC seed regions did not show any connection with the temporal and inferior parietal cortices. Overall, all DLPFC seeds showed strong connectivity with the FP, IFG, and the limbic system. The tractography results suggest a single axis of changing connectivity from ventral-caudal to dorsal-rostral regions, with the key differences being between the most ventral-caudal regions and elsewhere. Specifically, the ventral-caudal seeds (9/46va and 9/46vp) show widespread structural connectivity to frontal, limbic, sensorimotor, and superior parietal cortex.

**Table 1.**
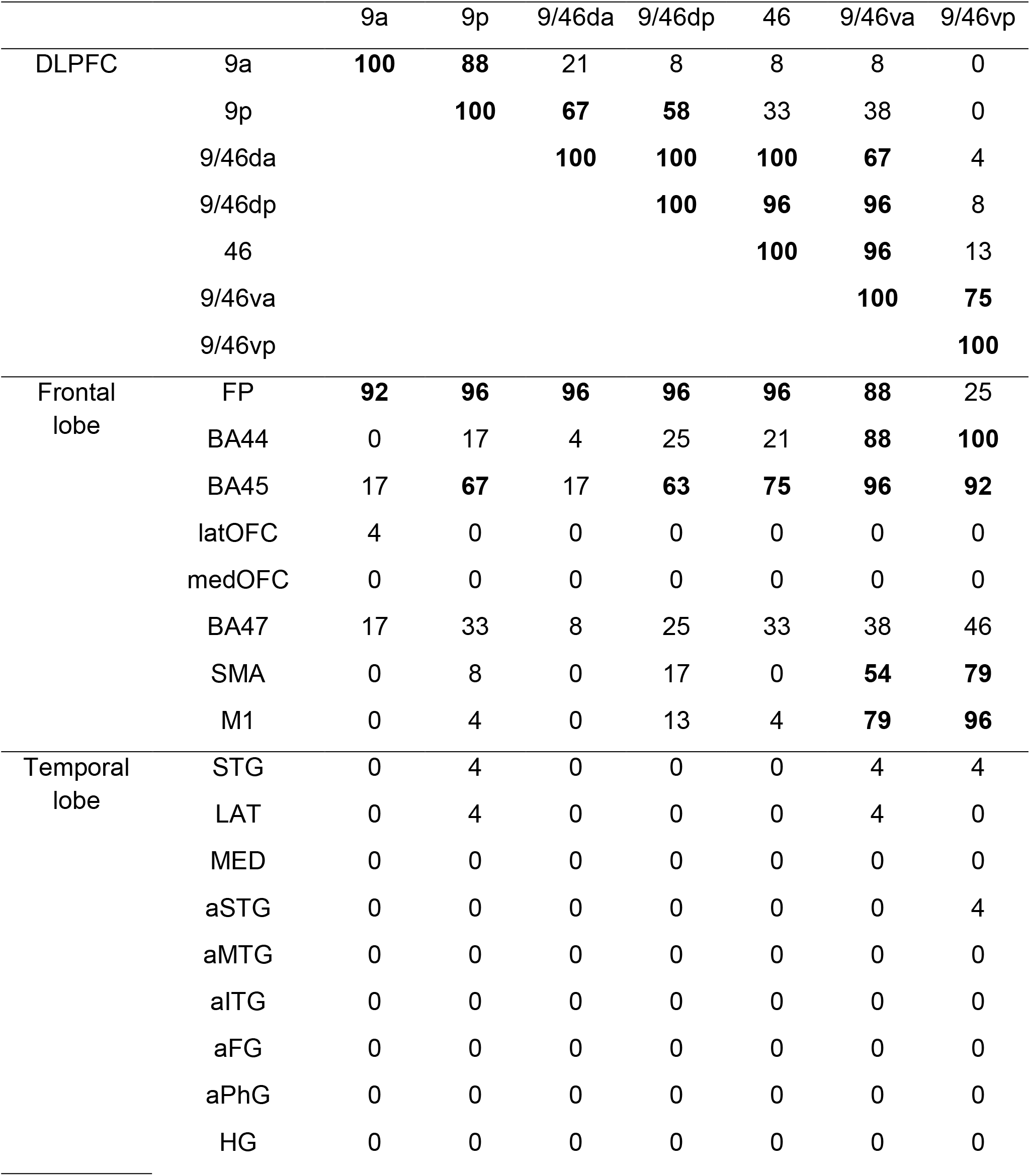
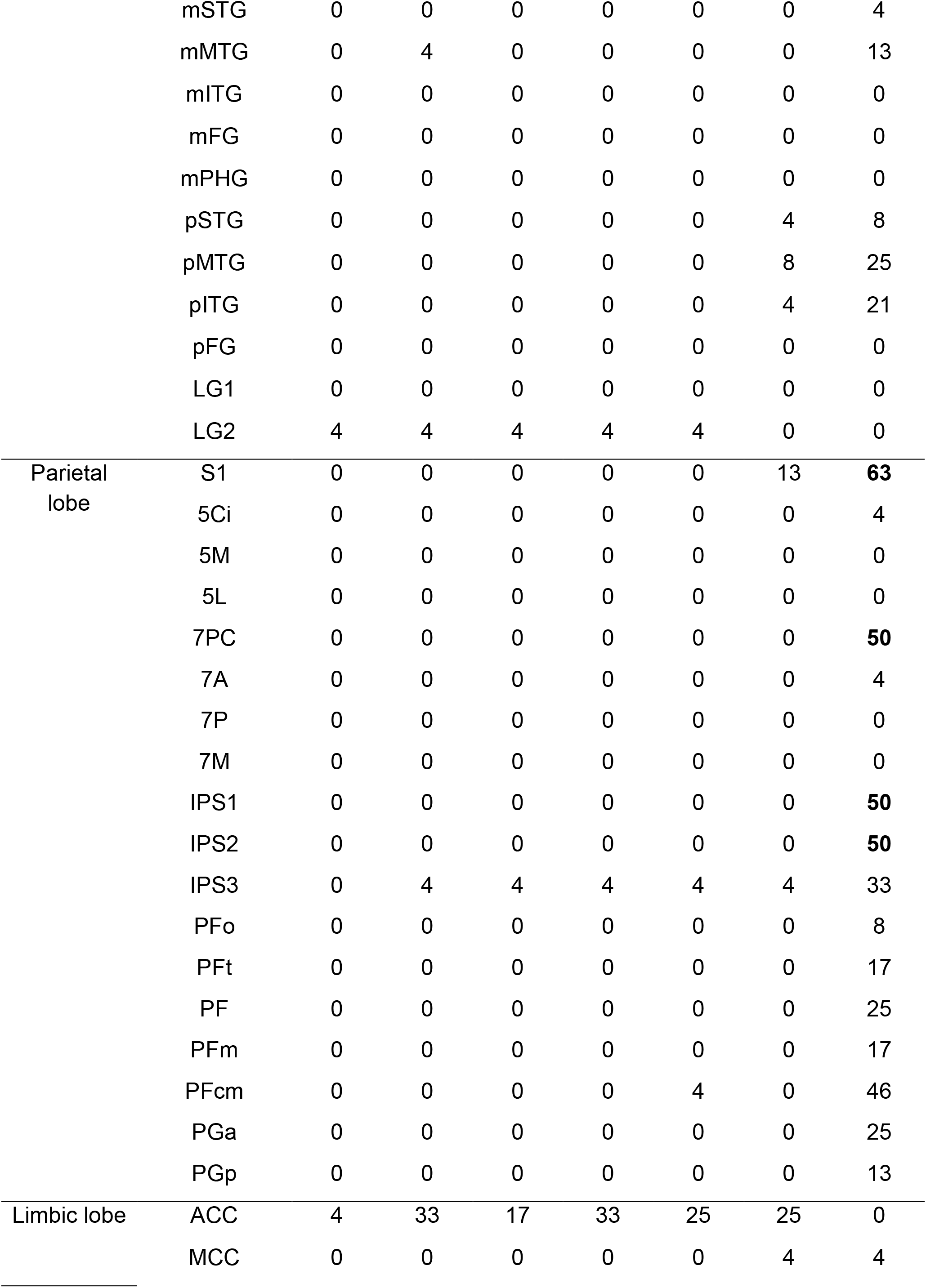
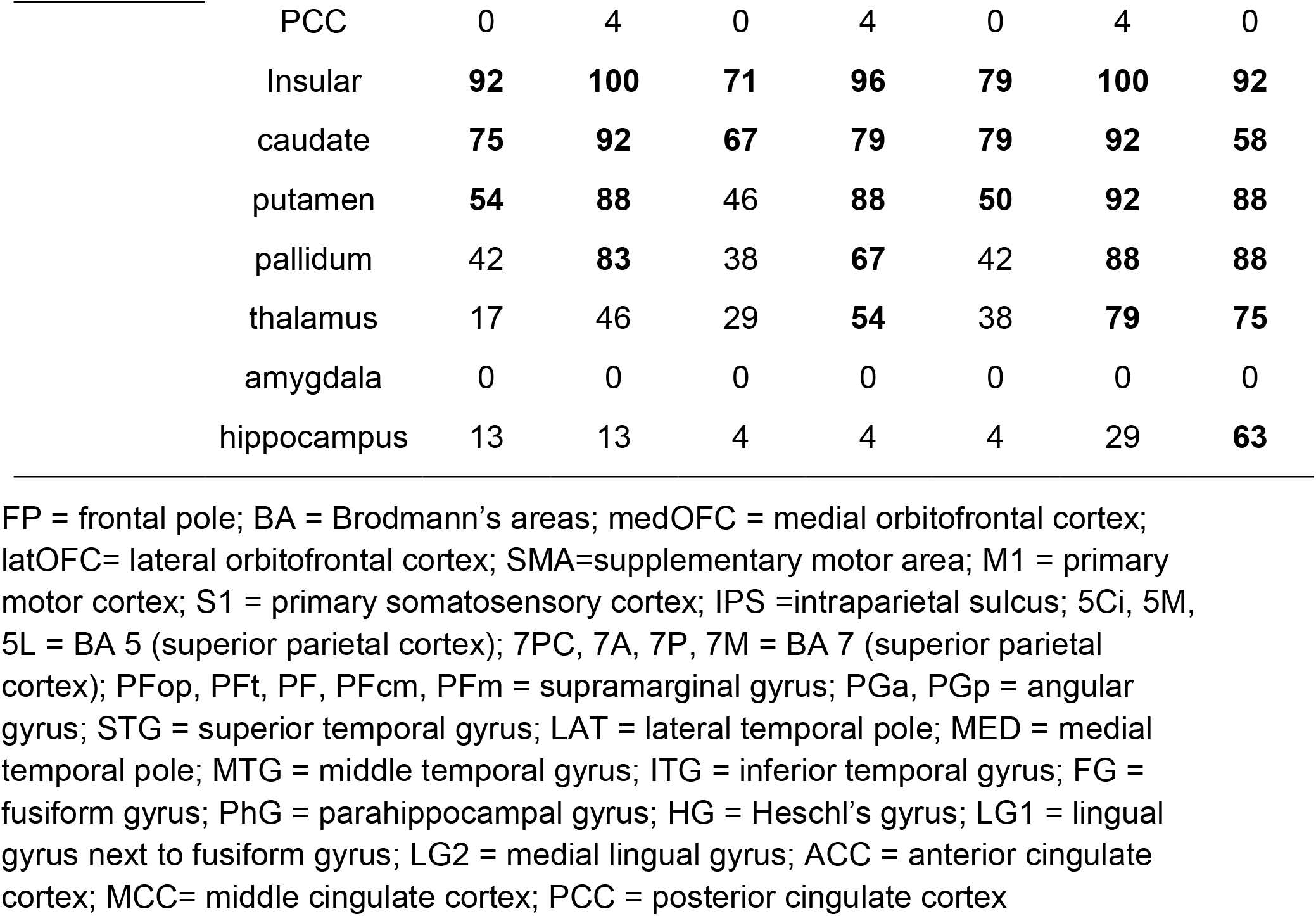
Structural connectivity results for each DLPFC region. Bold font indicates that the connection probability was over 50% (12/24) for group analysis.

**Figure 1.**
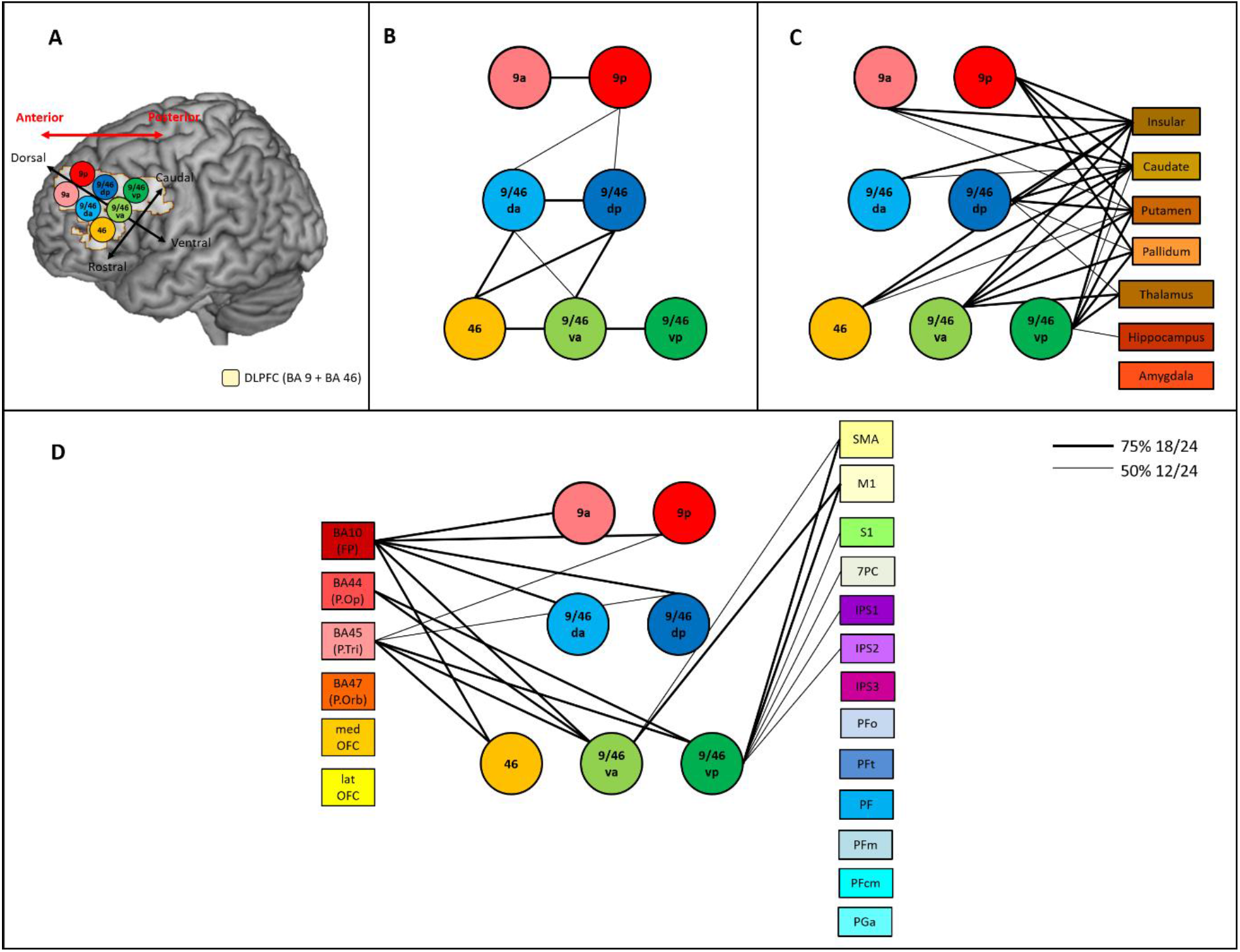
(A) The location of the seven DLPFC areas used as seed regions for the connectivity analyses. Red arrow indicate the anterior-posterior axis of the lateral prefrontal cortex. Black arrows represent each axis of the subregions of DLPFC. (B) Intra-DLPFC structural connectivity. (C) The structural connectivity between DLPFC seed regions and the limbic regions. (D) The structural connectivity between DLPFC seed regions and the frontal and parietal regions. Each DLPFC seed is represented by a circle. Lines connecting ROIs are displayed if the probabilistic tractography exceed the minimum probability threshold in either 50% (thin line) or 75% (thick line) of the participants.

**Figure 2.**
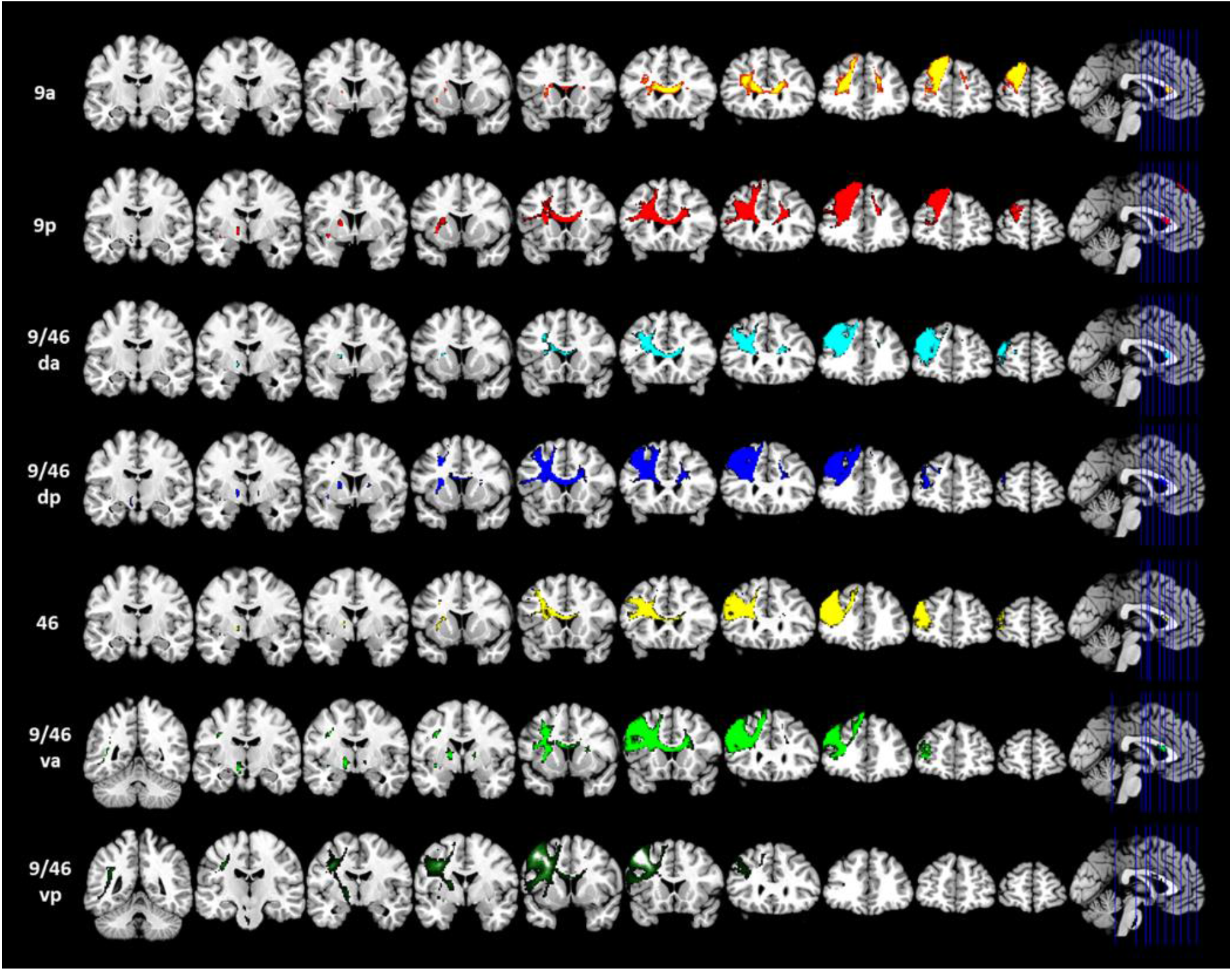
Structural connectivity patterns of the DLPFC seed regions.

FP = frontal pole; BA = Brodmann’s areas; medOFC = medial orbitofrontal cortex; latOFC= lateral orbitofrontal cortex; SMA=supplementary motor area; M1 = primary motor cortex; S1 = primary somatosensory cortex; IPS =intraparietal sulcus; 5Ci, 5M, 5L = BA 5 (superior parietal cortex); 7PC, 7A, 7P, 7M = BA 7 (superior parietal cortex); PFop, PFt, PF, PFcm, PFm = supramarginal gyrus; PGa, PGp = angular gyrus; STG = superior temporal gyrus; LAT = lateral temporal pole; MED = medial temporal pole; MTG = middle temporal gyrus; ITG = inferior temporal gyrus; FG = fusiform gyrus; PhG = parahippocampal gyrus; HG = Heschl’s gyrus; LG1 = lingual gyrus next to fusiform gyrus; LG2 = medial lingual gyrus; ACC = anterior cingulate cortex; MCC= middle cingulate cortex; PCC = posterior cingulate cortex

### Functional connectivity patterns across the DLPFC

The whole-brain resting-state functional connectivity (rsFC) map of each DLPFC seed region is displayed in Fig. 3. Overall, the 7 ROIs showed involvement in two distinct networks with graded rsFC patterns suggesting a transition between these networks. The two BA 9 seeds (9a and 9p) were primarily correlated with the regions of the default mode network (DMN) including medial prefrontal cortex (mPFC), OFC, IPC (particularly angular gyrus), precuneus, PCC, anterior/middle temporal regions, and hippocampus (Raichle et al., 2001; Buckner et al., 2008). All other seed regions were strongly correlated with brain regions of the multiple demanding network (MDN) including IFG, SMA, ACC/MCC, SPC, IPS, supramarginal gyrus, and pMTG (Duncan and Owen, 2000; Seeley et al., 2007; Woolgar et al., 2011; Spreng et al., 2013). However, area 9/46vp showed connectivity with both the MDN and the DMN. All DLPFC seed regions were strongly functionally connected to the insular and basal ganglia regions. The results look to vary along with dorsal-ventral axis such that the dorsal parts of the DLPFC are connected with the DMN, whereas the ventral parts of DLPFC are associated with the MDN. Similar to the structural connectivity results, the most ventral-caudal seed (9/46vp) shows the widespread functional connectivity across both of the DMN and MDN.

**Figure 3.**
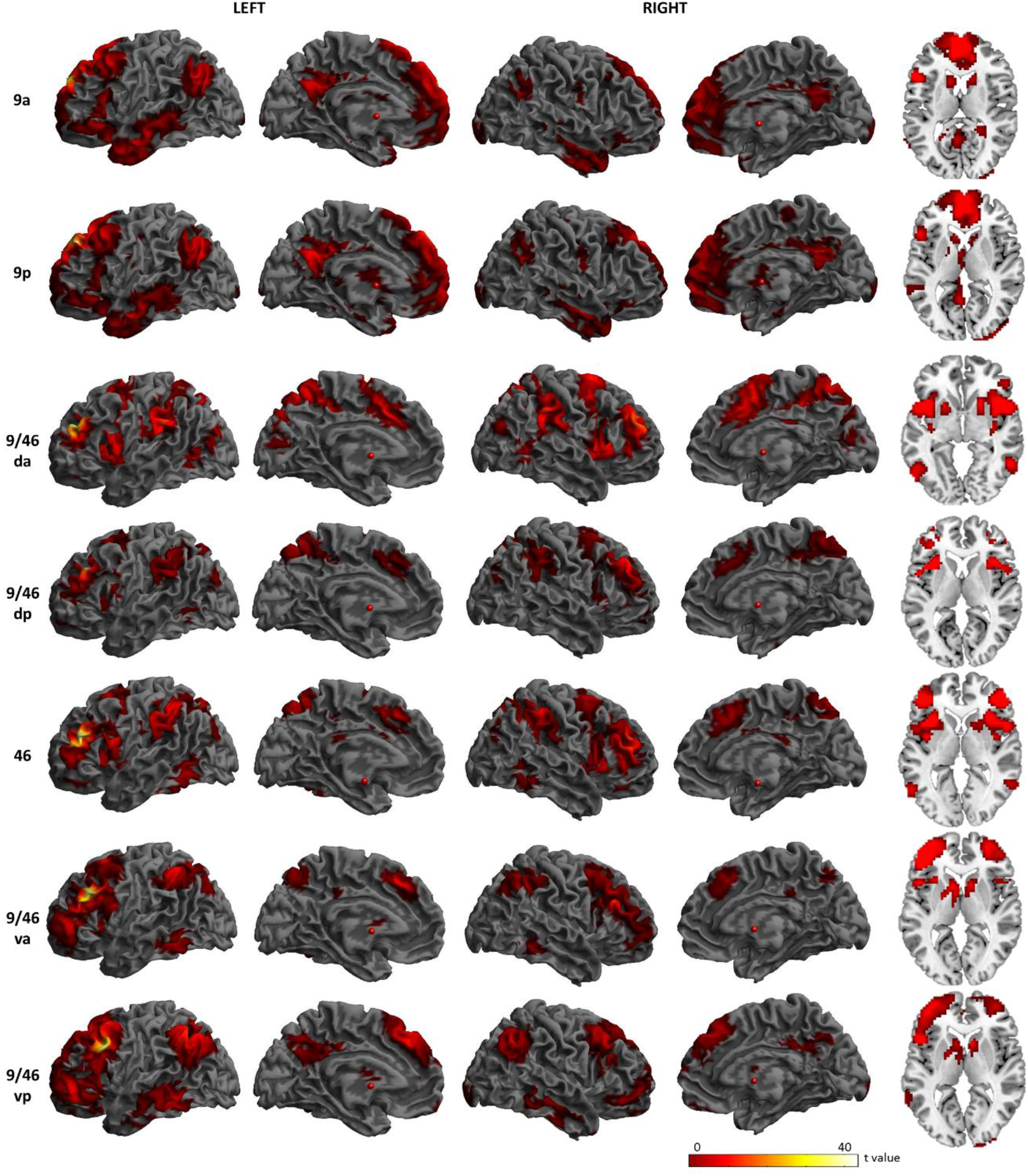
Functional connectivity patterns of the DLPFC seed regions.

To quantify the differences in rsFC across the DLPFC and visualise the shifting connectivity across the critical axes, the rsFC maps were compared between pairs of DLPFC seed regions varying along the rostral-caudal axis. Fig. 4 shows the result of comparisons within each gyrus, along the rostral-caudal axis. 9a revealed stronger rsFC with the insula and IPL than 9p, whereas 9p showed higher rsFC with mPFC, angular gyrus, and precuneus than 9a. 9/46da showed higher rsFC with the IFG, insula, M1/S1, MCC, SPL, IPL, ITG, and visual cortex, yet lower rsFC with IPC, precuneus, PCC, and lateral temporal cortex than 9/46dp. The comparisons between the more rostral and caudal ventral seed regions exhibited prominent differences in similar regions. Relatively rostral regions showed higher rsFC with regions of the MDN including the IFG, SMA, M1/S1, supramarginal gyrus, ACC/MCC and visual cortex, yet lower rsFC with DMN regions, such as the mPFC, OFC, angular gyrus, precuneus, PCC, and lateral temporal cortex, than more caudal regions.

**Figure 4.**
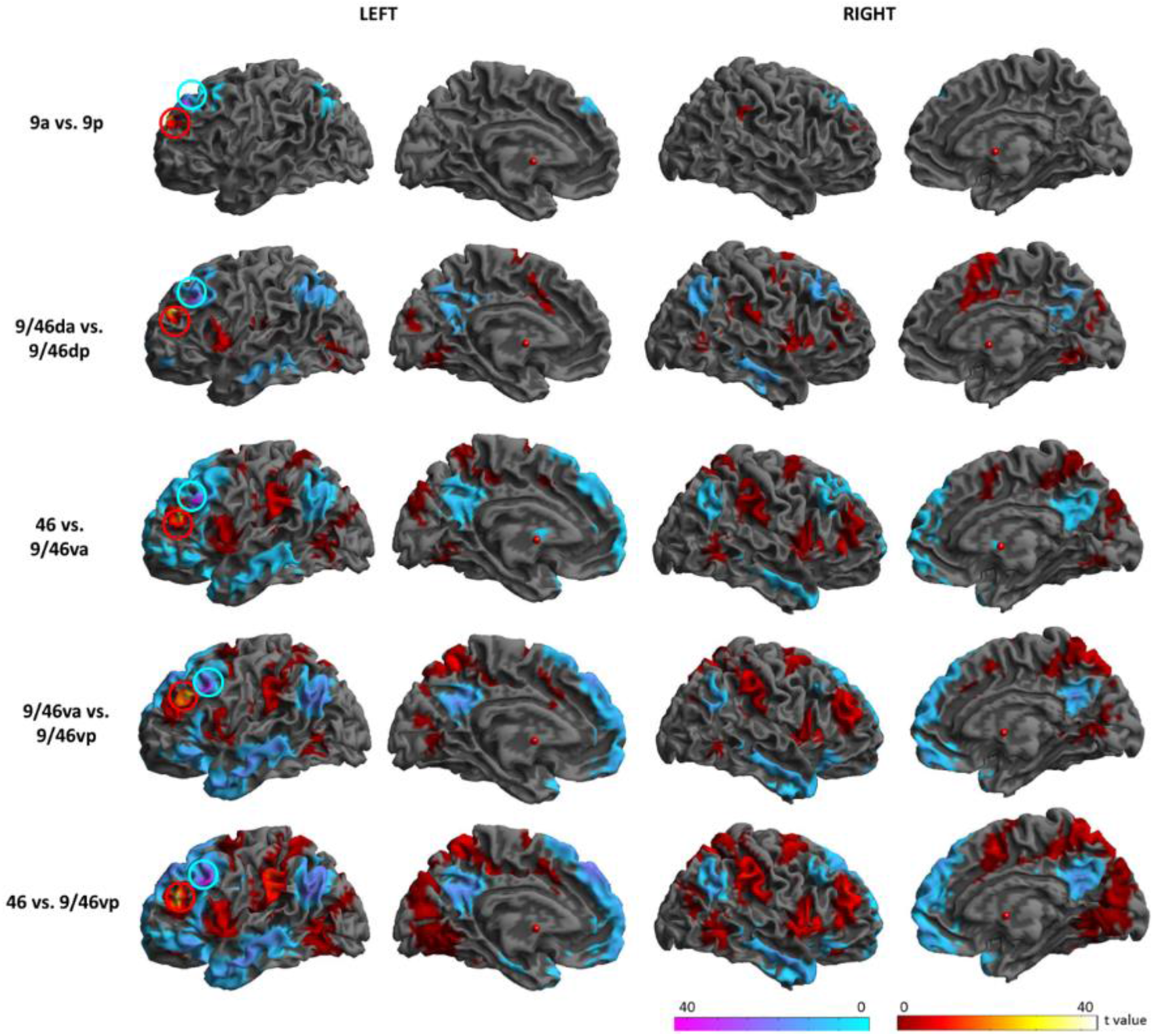
Comparison of the rsFC along the rostral-caudal axis. Circles indicate the DLPFC seed regions. Warm colours indicate the comparison from the rostral to the caudal regions. Cold colours indicate the comparison from the caudal to the rostral regions.

In order to compute the differences between seed regions along with dorsal-ventral axis, we combined the each set of seed regions on the rostral-caudal axis. Fig. 5 shows the result of comparisons along the dorsal-ventral axis. Dorsal regions (9a and 9p) had significantly higher rsFC with the regions in the DMN and lower rsFC with the parts of the MDN than the middle regions (9/46da and 9/46dp). The middle regions showed higher rsFC with the MDN, yet lower rsFC with the DMN than ventral regions (46, 9/46va, and 9/46vp). The ventral regions had significantly higher rsFC with the MDN, whereas lower rsFC with the DMN than the dorsal regions.

**Figure 5.**
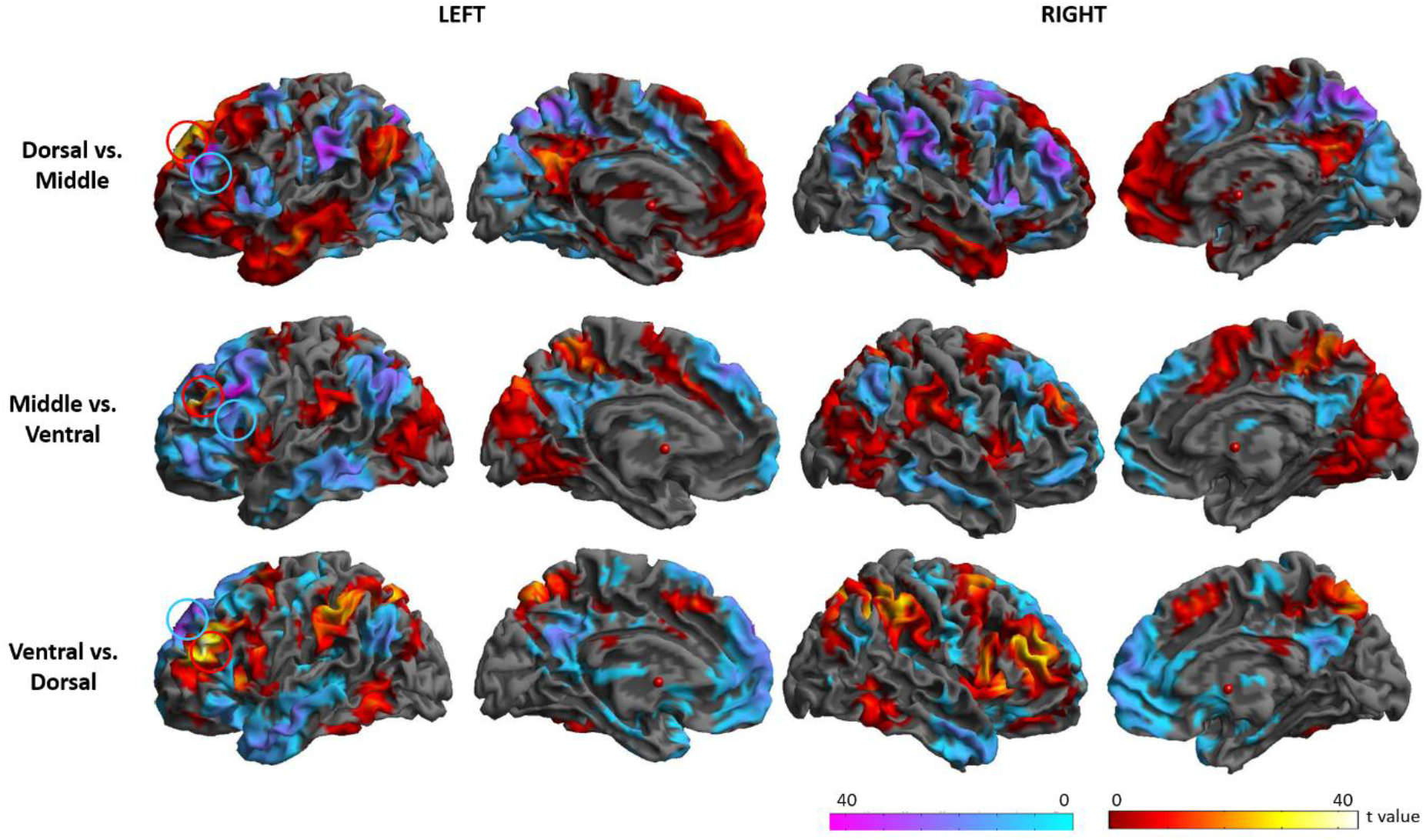
Comparison of the rsFC along the dorsal-ventral axis. Circles indicate the location of the DLPFC seed regions.

Overall, the dorsal parts of the DLPFC had strong connectivity with the DMN, whereas the ventral DLPFC regions were strongly connected with the MDN.

### Structural and functional connectivity profiles of DLPFC seed regions

The connectivity profile of the DLPFC seed regions is displayed in Fig. 6. Overall, the seed regions showed more widespread connections to target regions functionally than structurally (although the structural connectivity of 9/46vp was quite extensive), with structural connectivity mainly limited to the frontal and limbic cortex. With a more liberal threshold in structural connectivity (25% of the participants), the dorsal-caudal seeds (9p and 9/46dp) and the ventral-rostral seed (46) showed a connection with the ACC and the most ventral-caudal seed (9/46vp) had a connection to the posterior MTG and angular gyrus (Fig. 3). Functional profiles of the DLPFC revealed the distinctive connectivity patterns of the BA 9 region was strongly coupled with the DMN areas and that of the BA 9/46vp was connected to the DMN as well as MDN. The other regions in middle and ventral DLPFC (BA 9/46da, 9/46dp, 46, and 9/46va) has strong functional connectivity with region in the MDN.

**Figure 6.**
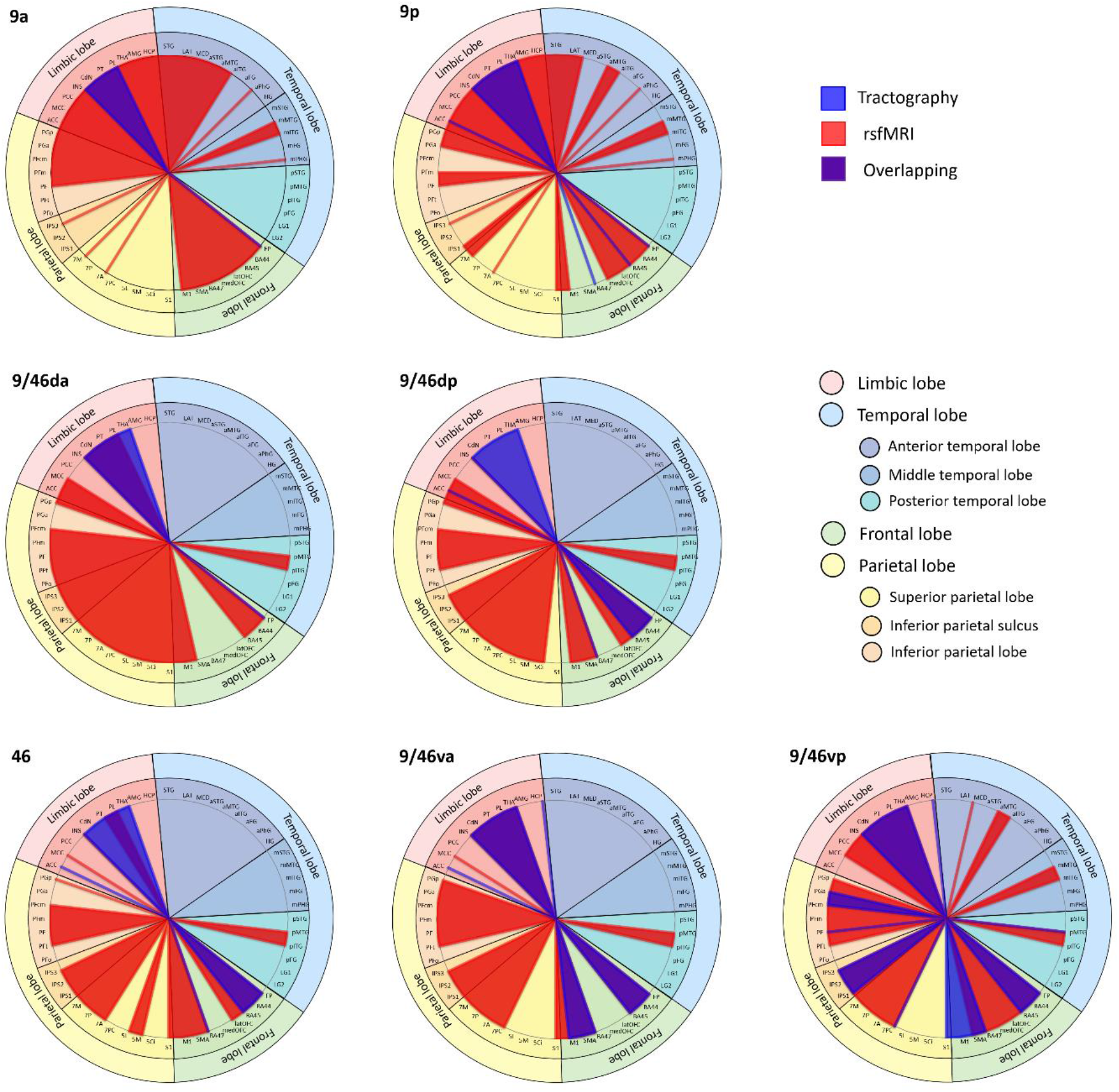
Summary of the structural (blue) and functional (red) connectivity profiles of the DLPFC seed regions. The overlapping areas are coloured in purple. The structural connectivity was thresholded at 25% of participants.

FP = frontal pole; BA = Brodmann’s areas; medOFC = medial orbitofrontal cortex; latOFC= lateral orbitofrontal cortex; SMA=supplementary motor area; M1 = primary motor cortex; S1 = primary somatosensory cortex; 5Ci, 5M, 5L = BA 5 (superior parietal cortex); 7PC, 7A, 7P, 7M = BA 7 (superior parietal cortex); IPS = inferior parietal sulcus; PFop, PFt, PF, PFcm, PFm = supramarginal gyrus; PGa, PGp = angular gyrus; STG = superior temporal gyrus; LAT = lateral temporal pole; MED = medial temporal pole; MTG = middle temporal gyrus; ITG = inferior temporal gyrus; FG = fusiform gyrus; PhG = parahippocampal gyrus; HG = Heschl’s gyrus; LG1 = lingual gyrus next to fusiform gyrus; LG2 = medial lingual gyrus; ACC = anterior cingulate cortex; MCC= middle cingulate cortex; PCC = posterior cingulate cortex; INS = insula; CdN = caudate nucleus; PT = putamen; PL = pallidum; THA = thalamus; AMG = amygdala; HCP = hippocampus

## Discussion

We investigated the patterns of connectivity in subregions of DLPFC along with the rostral-caudal and dorsal-ventral axes. We showed that subregions of DLPFC had differential structural and functional connectivity and distinctive patterns of DLPFC connectivity may be subdivided the DLPFC into subregions based on their structural and functional connectivity. Structural connectivity demonstrated graded intra-regional connectivity within the DLPFC. The patterns of connectivity between the DLPFC subregions and other cortical areas revealed a separation of dorsal-rostral subregions from the most ventral-caudal subregion. The dorsal-rostral subregions was restricted to link other frontal and limbic areas whereas the ventral-caudal region was widely connected to frontal, temporal, parietal, and limbic cortex. The patterns of functional connectivity revealed that subregions of DLPFC were strongly interconnected to each other within the whole frontal cortex and coupled with two functional brain networks: MDN and DMN. The dorsal subregions were associated with the DMN, while middle dorsal-rostral subregions were linked with the MDN, respectively. Similar to the results of structural connectivity, the most ventral-caudal subregion showed increased functional coupling with both DMN and MDN. Our results suggest that DLPFC may be subdivided by the diagonal axis of the dorsal-ventral axis and rostral-caudal axis. Our findings support the framework of a functional organization along the anterior-posterior axis in the lateral prefrontal cortex (Petrides and Pandya, 1999; Koechlin et al., 2003).

The Cascade model explains the process that executive control is implemented within the lateral prefrontal cortex along a posterior-to-anterior hierarchy, from simple to more abstract cognitive control processing (Koechlin et al., 2003). For example, posterior DLPFC supports action selection based on sensory input and anterior DLPFC provides episodic control for action selection, taking into account the ongoing context. The frontopolar cortex supports branching control for action selection based on a holding temporal context. Our structural connectivity results supports this progressive posterior to anterior hierarchy within the DLPFC subregions, showing highly interconnected subregions within each gyri via short U-fibres as well as DLPFC connections with the FP, IFG and motor regions via short frontal tracks (Catani et al., 2012; Yeterian et al., 2012). Specifically, the dorsal-rostral subregions connected to the FP via the frontal aslant tract, while the ventral-caudal subregions had connection to sensory motor regions through the frontal longitudinal tracts. Similarly, our functional connectivity results demonstrated that subregions of DLPFC had strong coupling with other frontal regions including the FP, IFG, OFC, and motor cortex. These patterns of connectivity within the DLPFC reflecting local short fibres suggest its graded and integrative organization, support the anterior-posterior gradient across the whole frontal system (Petrides and Pandya, 1999; Petrides, 2005b).

We observed that the dorsal-rostral subregions (anterior parts of the DLPFC) were linked to the frontopolar regions with increased functional connectivity with the DMN. As the frontopolar cortex is a supramodal area involved in various higher order functions such as self-directed thought, rational integration - the simultaneous consideration of multiple relations, and cognitive branching – holding goals while exploring secondary goals, planning, and reasoning (Ramnani and Owen, 2004). With co-activation of the frontopolar cortex, it has reported that the DMN could be activated for self-generated thought (Christoff et al., 2016) or increased cognitive reasoning complexity (Sormaz et al., 2018). These studies supports our findings that dorsal-rostral subregions of the DLPFC connected with the FP were strongly coupled with the DMN. In line with the anterior-posterior gradient in the prefrontal cortex, our connectivity analysis suggests that the anterior parts of the DLPFC would be involved in more challenging cognitive control such as complex cognitive reasoning and cognitive branching.

In contrast, the middle-ventral subregions (middle-posterior parts) were connected to the IFG (BA 44 and 45) with strong coupling with the MDN and the ventral-caudal region (the most posterior part) had anatomical connection with temporal and parietal areas with increased functional connectivity with both DMN and MDN. Several cortico-cortical association pathways link the prefrontal cortex and other cortical regions (Petrides and Pandya, 1984; Petrides, 2005b; Catani et al., 2012; Thiebaut de Schotten et al., 2012; Yeterian et al., 2012). The superior longitudinal fasciculus (SLF) links the PFC and parietal cortex. SLF has three distinct branches: SLF I connecting the superior frontal area (BA 8, 9, 32) to SPC, SLF II connecting the SFG/MFG to IPS/AG, SLF III connecting IFG to IPS. The arcuate fasciculus (AF) connects the posterior regions of the frontal lobe and temporal lobe (Parker et al., 2005). The inferior fronto-occipital fasciculus (IFOF) connects occipital cortex, temporal areas, ventrolateral frontal cortex and inferior parietal regions (Schmahmann et al., 2007; Martino et al., 2010). As a part of the MDN, the IFG is involved in cognitive control and language processing (Brass et al., 2005; Camilleri et al., 2018). As the IPS shows anatomical connection with the DLPFC via SLF I/SLF II (Petrides and Pandya, 1984; Petrides, 2005a; Thiebaut de Schotten et al., 2012), the IPS acts as a multifaceted behavioural integrator, binding task-relevant information from the sensory, motor, and cognitive domains, mediated by the top-down control of DLPFC (Gottlieb, 2007). These findings suggest that the posterior parts of the DLPFC would be associated with the core processes of cognitive control, supporting the anterior-to-posterior functional organization of the DLPFC.

In our results, the middle-ventral subregions did not show anatomical connections with the IPS but they were functionally coupled with the MDN. One explanation of this discrepancy is that a weak anatomical connection between two regions may still hold a high functional significance via indirect connections of shared brain regions (Friston, 2002; Damoiseaux and Greicius, 2009). Functional connectivity does not necessarily require direct, physical connections and several studies have reported functional connections between regions without anatomical connectivity (Damoiseaux and Greicius, 2009). Therefore, the functional connectivity without physical connections potentially results from indirect anatomical connections via shared brain areas.

The DLPFC is functionally and structurally connected with subcortical areas including the insular and ACC (Catani et al., 2012; Cieslik et al., 2013). As core areas of the MDN, insular and ACC play a role in cognitively demanding tasks, responding to uncertainty and emotional salience (Seeley et al., 2007; Menon and Uddin, 2010; Camilleri et al., 2018). A meta-analysis study demonstrated strong functional connectivity between DLPFC and insular/ACC (Cieslik et al., 2013). We also showed significant functional connectivity between insular/ACC and the DLPFC. However, our tractography showed structural connections between the DLPFC and insular only, not the ACC. With a lower threshold, we found some evidence of a connection between the DLPFC regions and ACC (25% of participants). In addition, we showed that the corticostriatum projections directly link all DLPFC subregions to the basal ganglia and thalamus (Alexander et al., 1986; Jarbo and Verstynen, 2015). In particular, the anatomical connections between the basal ganglia and DLPFC form a neural circuit involved in several aspects of goal directed behaviours (for a review, see Haber, 2003), which supports a role for the DLPFC in action control (Petrides, 2005b; Cieslik et al., 2013). Furthermore, the extensive connections from the basal ganglia to the cerebral cortex potentially account for the discrepancy between the structural and functional connectivity in the DLPFC subregions.

In the current study, we explored the structural and functional connectivity across the subregions of DLPFC using probabilistic tractography and rsfMRI approaches. The key limitations of the probabilistic tractography are the issues of distance effect and thresholding (Jones, 2008; Morris et al., 2008). A degree of uncertainty in fibre orientation exists at each step in the propagation of a pathway. This accumulation of uncertainty from voxel to voxel as the streamline is advanced causes a decrease in probability with increasing path length and a progressive dispersion of the streamlines with distance from the seed (Morris et al., 2008). Therefore, it is difficult to determine a threshold value which will identify true positives while simultaneously minimising the rate of both Type I errors in regions close to the seed and Type II errors in distant regions. Although our procedure most likely produced a conservative cut-off value for longer pathways (Binney et al., 2012; Cloutman et al., 2012), there may be long-range connections left undetected.

## Acknowledgments

This research was supported by a Beacon Anne McLaren Research Fellowship (University of Nottingham) to JJ, a British Academy Postdoctoral Fellowship awarded to RLJ (pf170068), and an Advanced ERC award (GAP: 670428 - BRAIN2MIND_NEUROCOMP), MRC programme grant (MR/R023883/1), and intramural funding (MC_UU_00005/18) to MALR.

## Author contribution

J.J: Conceptualization, Methodology, Formal analysis, Investigation, Data curation, Writing - original draft, Writing - review & editing, Visualization.

M.A.L.R: Methodology, Investigation, Writing - review & editing.

R.L.J: Methodology, Writing - original draft, Writing - review & editing.

